# Distinct RanBP1 nuclear export and cargo dissociation mechanisms between fungi and animals

**DOI:** 10.1101/294686

**Authors:** Yuling Li, Jinhan Zhou, Yuqing Zhang, Qiao Zhou, Xiaofei Shen, Da Jia, Junhong Han, Qingxiang Sun

## Abstract

Ran binding protein 1 (RanBP1), the primary effector of nuclear GTPase Ran, is a cytoplasmic-enriched and nuclear-cytoplasmic shuttling protein, playing important roles in nuclear transport through preventing RanGTP from being trapped with karyopherin proteins and dissociating cargoes from nuclear export factor CRM1. Much of what we know about RanBP1 is learned from fungi. Here we show that animal RanBP1 has distinct cargo dissociation and nuclear export mechanisms. In contrast to CRM1-RanGTP sequestration mechanism of cargo dissociation in fungi, animal RanBP1 solely sequesters RanGTP from nuclear export complexes. In fungi, RanBP1, CRM1 and RanGTP form a 1:1:1 nuclear export complex; in contrast, animal RanBP1, CRM1 and RanGTP form a 1:1:2 nuclear export complex. The key feature for the two mechanistic changes from fungi to animals is the loss of affinity between RanBP1-RanGTP and CRM1, since residues mediating their interaction in fungi are not conserved in animals. The biological significances of these different mechanisms in fungi and animals are also studied and discussed. Our study illustrates how orthologous proteins may play conserved functions through distinct routes, and may provide directions for design of antifungal medicines.

## Introduction

Eukaryotic cells have nucleus which segregates the nucleoplasm and cytoplasm into two isolated compartments. Exchange between these compartments are mainly mediated through nuclear pore complex (NPC), a semi-permeable channel that allows only certain classes of molecules to pass through, e.g. importins and exportins, which are collectively called karyopherin proteins ^1^. Cargo entering nucleus must possess a nuclear localization signal (NLS), which binds to an importin and enters nucleus though NPC ^2^. In the nucleus, the GTP-bound form of Ran (Ras-related nuclear) protein dissociates the importin-NLS cargo and RanGTP-Importin is exported to cytoplasm ^3,4^. For cargo’s nuclear export, its nuclear export signal (NES) forms a complex with exportin in the presence of RanGTP and together the trimeric complex translocates to the cytoplasm through NPC ^5^. In the cytoplasm, RanGTP complexes (with either importin or exportin-NES) are hydrolysed to RanGDP by GTPase activating protein RanGAP, dissembling the complexes and recycling karyopherins and Ran for further rounds of nuclear transport ^6^.

However, importin or exportin-NES displays extremely high affinity (in the nM range) for RanGTP and inhibits RanGAP-facilitated RanGTP hydrolysis ^7,8^. Efficient hydrolysis requires Ran binding proteins containing one or more Ran binding domains (RBDs, around 150 residues each), to dissociate RanGTP from karyopherin prior to hydrolysis ^9–11^. In human, there are two such proteins, namely RanBP1 and RanBP2. While the predominantly cytoplasmic RanBP1 contains one RBD, the cytoplasmic rim-attached RanBP2 has four RBDs^10^. These RBDs are the tightest binders of RanGTP, K_d_ being around 1 nM, whereas it binds to RanGDP at only approximately 10 µM affinity ^12–14^.

In addition, RanBP1 in the cytoplasm functions in the disassembly of nuclear export complexes (also before RanGTP is hydrolysed), through dissociating NES containing cargoes from the complexes ^15^. CRM1 (Chromosomal Region Maintenance 1, also known as Exportin-1) is a major nuclear export factor that is responsible for nuclear export of a plethora of proteins containing NES sequence(s) ^16^. In yeast, after binding to RanGTP and CRM1,RanBP1 allosterically closes the groove, releasing CRM1 cargoes into the cytoplasm ^17^.

Sequence analysis indicates that in contrast with fungi RanBP1 (which includes yeast RanBP1), animal RanBP1 contains NES sequence C-terminal to its RBD ^18^ (Fig. S1). It is reported that the NES of human RanBP1 (hRanBP1) is responsible for its cytoplasmic accumulation (RanBP1 is a shuttling protein) ^18,19^. If animal RanBP1 binds to RanGTP-CRM1 similarly as yeast RanBP1 (yRanBP1), an apparent paradox then exists: its NES binding to CRM1 is inhibited by its own RBD. Specifically, if animal RanBP1 binds CRM1 through NES in the nucleus to prepare for nuclear export, its RBD might immediately interact with RanGTP on CRM1 (because of proximity and high affinity) and dissociate its own NES before animal RanBP1 is exported. One may argue that NES may play a role in recruiting animal RanBP1 to CRM1. However, RanBP1-RanGTP-CRM1 complex displays much higher affinity than NES- CRM1-RanGTP complex in yeast ^20^, thus the recruiting purpose seems unnecessary and unlikely. It should be noted that there are about ten residues between RBD and NES, which is insufficient to cover the distance (about 70 Å) between RBD and NES in space. Therefore in theory, the NES and RBD of animal RanBP1 would not bind to CRM1-RanGTP simultaneously. It is fascinating as to why animal RanBP1 requires an extra NES while fungi RanBP1 does not; what factor(s) prevents animal RBD from dissociating its own NES during its nuclear export; whether the NES of animal RanBP1 functions in cargo dissociation by direct competition with NES of cargo; and how animal RBD and NES binding to CRM1- RanGTP is regulated in time and space in cells. Intrigued by these long-standing questions, we performed biochemical, biophysical and cellular studies on RanBP1 and related proteins. Our work not only solved those puzzles, but also discovered unexpected animal RanBP1 nuclear export and cargo dissociation mechanisms distinctive from those in the yeast.

## Materials and Methods

### Cloning, protein expression and purification

The human, mouse or yeast RanBP1 (or their mutants) was cloned into a pGEX-4T1 based expression vector incorporating a TEV-cleavable N-terminal GST-tag fusion. The plasmid was transformed into Escherichia coli BL-21 (DE3) and grown in LB Broth medium. Expression of protein was induced by the addition of 0.5 mM isopropyl β-D-1-thiogalactopyranoside (IPTG) and the culture was grown overnight at 18˚C. Cells were harvested and sonicated in lysis buffer (50 mM Tris pH 8.0, 200 mM NaCl, 10% glycerol, 2 mM DTT, 1 mM EDTA and 1 mM PMSF). RanBP1 was purified on a GST column and eluted after TEV cleavage in buffer containing 20 mM Tris pH 8.0, 200 mM NaCl, 10% glycerol, 1 mM EDTA and 2 mM DTT followed by a Superdex 200 increase gel filtration column on the ÄktaPure (GE Healthcare) using the gel filtration buffer (20 mM Tris pH 8.0, 200 mM NaCl, 10% glycerol, 2 mM DTT). Eluted proteins were frozen at −80 °C at 5-10 mg/ml. Alternatively, GST-BP1 was eluted with 20 mM Tris pH 8.0, 200 mM NaCl, 1 mM EDTA, 2 mM DTT, 10 mM reduced glutathione and purified by Superdex 200 increase column. His-tagged proteins (human and yeast CRM1 and related mutants) were expressed in *E.coli* grown in TB Broth medium. The proteins were induced in the presence of 0.5 mM IPTG overnight at 25˚C, and purified by Nickel beads. 6× His-tagged proteins were eluted with 300 mM Imidazole pH 7.5, 300 mM NaCl, 10% glycerol and 2 mM BME.

### Pull down assay

All proteins used are purified by S200 prior to pull down. To assess different interactions, GST-tagged proteins were immobilized on GSH beads. For GST-RanBP1, a wash step was performed immediately after immobilization to remove unbound GST-RanBP1. Soluble proteins at indicated concentrations were incubated with the immobilized proteins in a total volume of 1ml for two hours at 4 °C. After two washing steps, bound proteins were separated by SDS/PAGE and visualized by Coomassie Blue staining. Each experiment was repeated at least twice and checked for consistency. Pull down buffer contains 20 mM Tris pH 8.0, 200 mM NaCl, 10% glycerol, 5 mM MgCl_2_, 0.005% Triton-X100 and 5 mM DTT if not specified.

### Isothermal Titration Calorimetry (ITC)

ITC experiments were conducted at 20 °C using ITC200 (Microcal) in buffer containing 20 mM Tris pH 8.0, 200 mM NaCl and 5 mM MgCl_2_. Q69L and L182A double mutant of Ran were used in this assay because this Ran mutant is 100% GTP bound (refer to co-submitted paper). For animal complex, 250 µM Ran was titrated into the sample cell containing 20 µM hCRM1 and 20 µM mRanBP1. For yeast complex, 150 µM Ran was titrated into the sample cell containing 15 µM yCRM1 and 15 µM yRanBP1. Each experiment was repeated at least twice. Data were processed by NITPIC ^21^ and fitted by SEDPHAT ^22^.

### Cell culture and transfection

HeLa cells were maintained in Dulbecco’s modified Eagles medium (Hyclone) supplemented with 10% (vol/vol) fetal bovine serum (Biological Industries), and transfected with TurboFect transfection reagent (Thermo Scientific). Images were acquired by Olympus FV-1000 confocal microscope, and were analysed using NIH ImageJ software.

### GAP hydrolysis assay

Human (0.4 µM) or yeast Ran (0.3 µM) protein was first mixed with hRanBP1 (1 µM) or yRanBP1 (1 µM) respectively. Then samples were briefly incubated with or without 1 µM CRM1 from respective species. Since Ran has very slow intrinsic nucleotide exchange rate, 0.5 µM RCC1 and 100 µM of GTP was added to reload Ran with GTP after each cycle of GAP mediated hydrolysis in all samples. The samples were then incubated with 0.3 µM yeast RanGAP for 0, 15, 45 and 75 minutes. The amount of free phosphate generated was measured using GTPase assay kit (Bioassay) and Multiskan FC microplate reader (Thermo Scientific). Each reaction was repeated four times in parallel.

## Results

### Mouse and yeast RanBP1 bind to CRM1-RanGTP differently

In yeast, RanBP1, RanGTP and CRM1 form a tight complex whereby RanBP1 forces H9 loop of CRM1 to allosterically close NES binding groove and dissociate NES. In order to visualize the mode of animal RanBP1’s binding to RanGTP-CRM1, we first attempted to solve the crystal structure of complex formed by mouse RanBP1 (mRanBP1), human RanGTP (hRanGTP) and human CRM1 (hCRM1). Both mouse and human RanBP1 have been used in this study and they are highly similar (95% identity, Fig. S2). Though the yeast complex was formed readily as expected, the equivalent complex with animal proteins was hardly formed under similar conditions (Fig. 1A). It should be noted that human and yeast Ran protein shares 83% sequence identity and are often used interchangeably ^17,23^. Further, human Ran^L182A^ used in this paper is loaded with approximately 80% of GTP (refer to co-submitted manuscript) and is often referred as RanGTP. When RanBP1, RanGTP and CRM1 were mixed at 5:3:1 molar ratio and passed through size exclusion chromatography column, the yeast proteins formed stable complex and were co-eluted, but not the case for the animal proteins (Fig. S3). Interestingly, greater amount of animal complex was formed when concentration of RanGTP is increased (Fig. 1B). In contrast, the yeast complex was not affected by RanGTP concentration (Fig. 1B). These results suggest that mRanBP1 is somewhat different from yRanBP1, in forming complex with RanGTP-CRM1.

**Figure 1.**
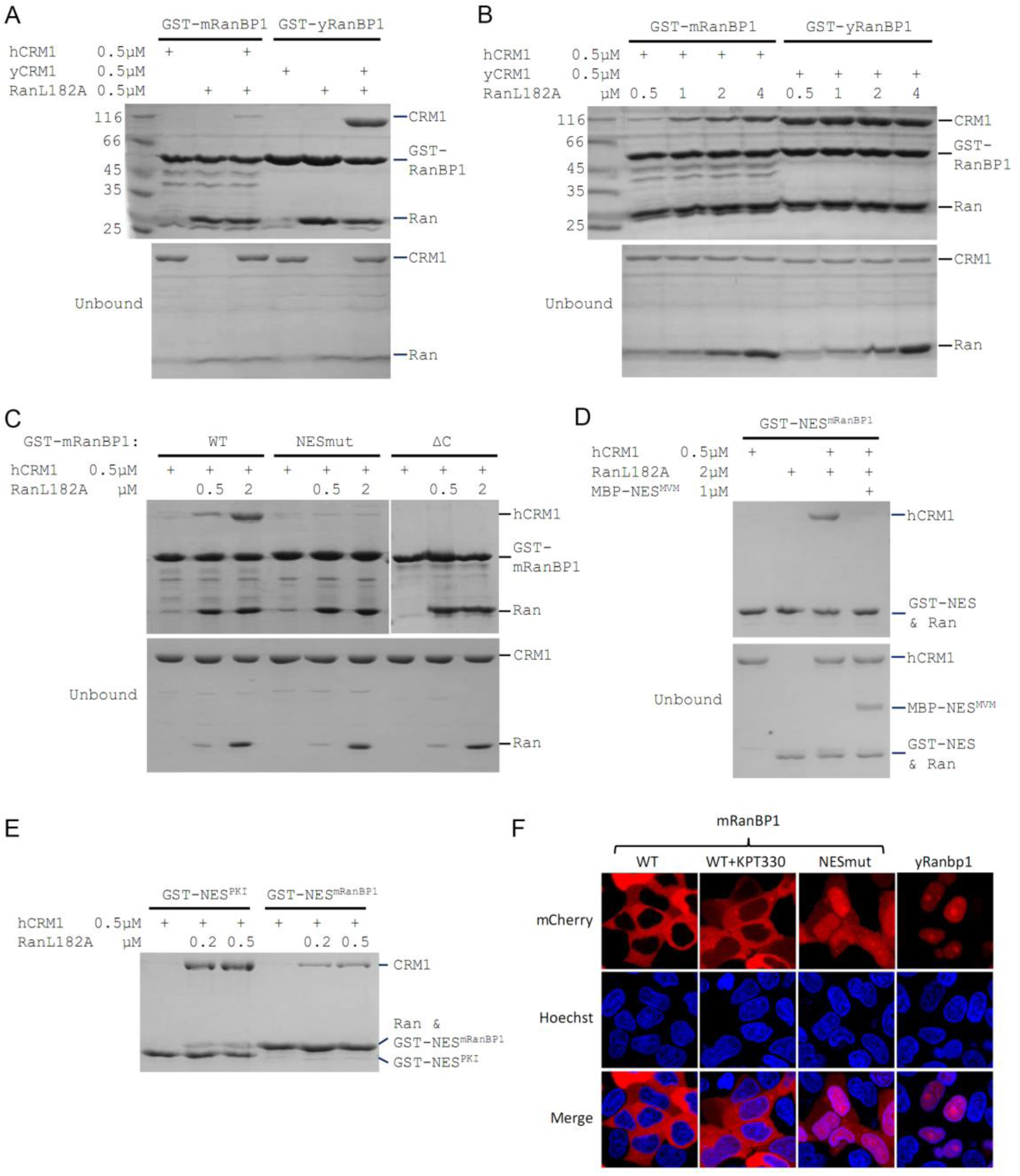
Mouse RanBP1 forms a nuclear export competent complex with hCRM1 only through its NES, and in a Ran concentration dependant manner. **(A, B, C)** GST tagged mouse/yeast RanBP1 or mutants pull down of CRM1 in the presence of RanGTP. **A)** While yeast proteins form a strong RanBP1-RanGTP-CRM1 complex, the animal proteins form a RanBP1-RanGTP complex with very weakly bound CRM1. **B)** At increasing concentration of RanGTP, while yeast complex is not affected, animal CRM1 is increasingly bound to GST- mRanBP1-RanGTP. **C)** Mutation (NESmut) or deletion of the NES of mRanBP1 (ΔC) abolishes hCRM1 binding at either high or low RanGTP concentration. **D)** GST-NES^mRanBP1^ pull down of CRM1 in the presence of RanGTP and NES inhibitor MBP-NES^MVM^. NES of mRanBP1 binds to hCRM1 in the presence of RanGTP and is effectively outcompeted by MBP-NES^MVM^, suggesting that the NES of mRanBP1 binds to NES groove of CRM1. **E)** GST-NES pull down of hCRM1 and different concentration of RanGTP to show that the NES of mRanBP1 is weaker than the well-studied NES of PKI cargo. **F)** mCherry tagged different RanBP1 constructs were transfected into HeLa cells and treated with or without 5 µM CRM1 inhibitor KPT-330 for 3h. While mRanBP1 is exclusively cytoplasmic, mRanBP1 treatment with KPT-330 and NESmut are significantly re-localized to the nucleus. yRanBP1 is exclusively nuclear.

### Mouse RanBP1’s NES is necessary for CRM1 binding and its nuclear export

To identify which region of mRanBP1 mediated the interaction with hCRM1, we cloned GST-tagged mRanBP1 mutants with NES mutation (termed as NESmut), the C terminus deletion (ΔC) or only the NES (NES^mRanBP1^). Surprisingly, both NESmut and ΔC lost binding to hCRM1 completely, in either low or high RanGTP concentration (Fig. 1C). In contrast, NES^mRanBP1^ bound to hCRM1 in the presence of RanGTP, and was outcompeted by supraphysiological NES ^24^ from minute virus of mice (Fig. 1D), suggesting that NES^mRanBP1^ is a regular NES that binds to NES groove on hCRM1. Further, the NES of mRanBP1 is weaker than that of regular NES from PKI (Protein kinase inhibitor) (Fig. 1E). Previously, it was reported that the NES of mRanBP1 was essential for its nuclear export ^25^. Indeed, while mRanBP1 was localized to the cytoplasm, NESmut (or mRanBP1 in the presence of CRM1 inhibitor KPT-330) was significantly re-localized to the nucleus (Fig. 1F). Unexpectedly, yRanBP1 was exclusively localized in the nucleus (Fig. 1F, right column), though RanBP1 proteins are known to be cytoplasmic localized proteins ^19,25^. The reason for this counter-intuitive observation will be explained in later sections. Altogether, these results conclude that the NES of mRanBP1 is necessary and sufficient for its interaction with hCRM1 in the presence of RanGTP, and this interaction is crucial for nuclear export of mRanBP1.

### Mouse RBD dissociates cargo through sequestering RanGTP

Previously we showed that ΔC or NESmut did not bind to hCRM1 in the presence of RanGTP (Fig. 1C). Relating to RanBP1’s function of CRM1 cargo dissociation, we wondered whether ΔC or NESmut possess cargo dissociation ability and whether NES^mRanBP1^ is involved in dissociating cargo, possibly through direct competition with cargo’s NES. Pull down assay showed that ΔC or NESmut dissociated cargoes as potently as WT mRanBP1 (Fig. 2A). This also means that NES of mRanBP1 is dispensable for cargo dissociation. Combined with the knowledge that the NES of mRanBP1 is weaker than the regular NES of cargo PKI (Fig. 1E), we believe that mRanBP1 does not dissociate cargo through direct competition of cargo’s NES.

**Figure 2.**
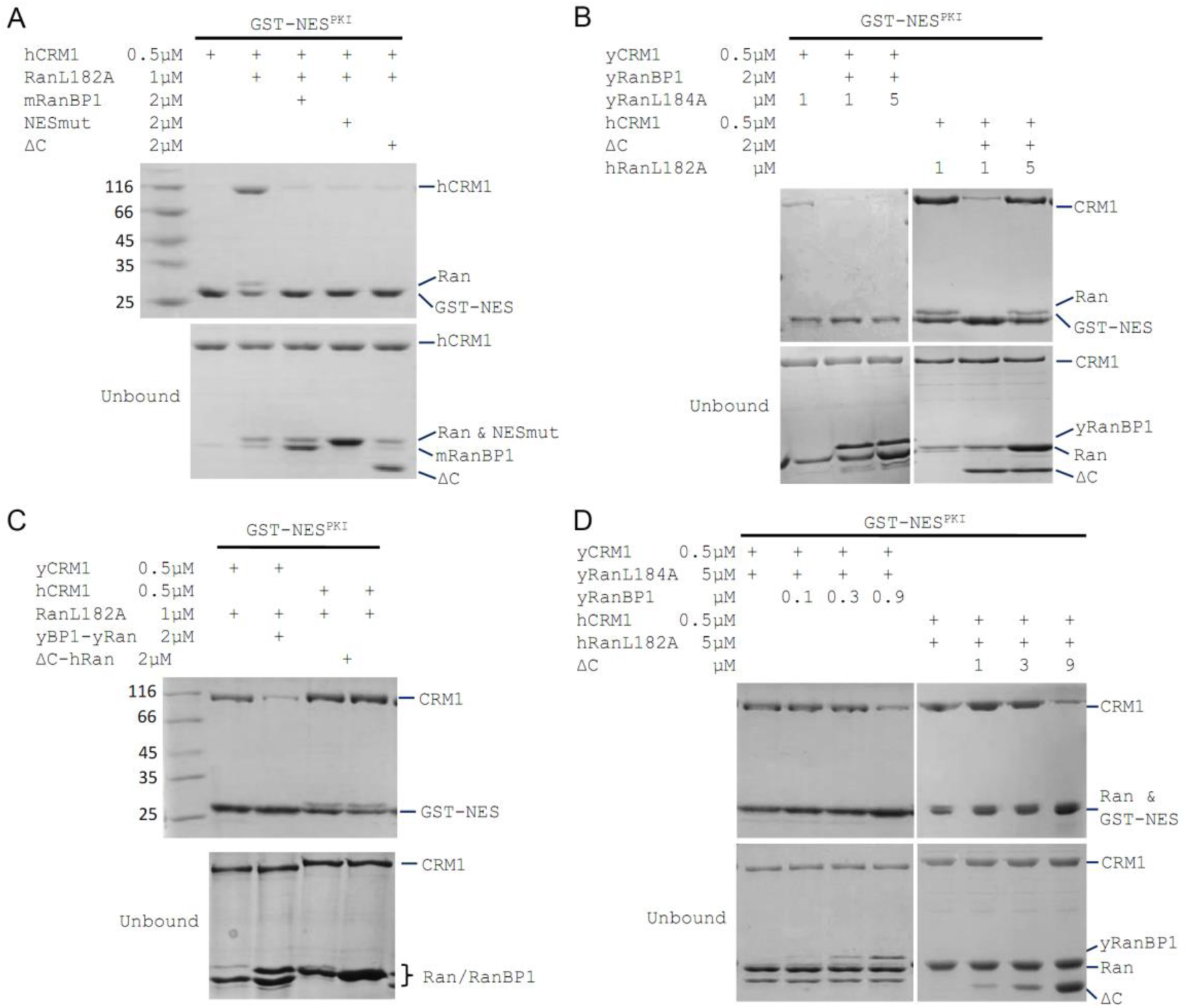
mRanBP1’s RBD dissociates CRM1 cargo through sequestering RanGTP. **(A-D)** GST- NES^PKI^ pull down of CRM1 and RanGTP in the presence of RanBP1 or its mutants. **A)** NESmut or ΔC dissociates cargo as potent as WT RanBP1. **B)** High concentration of RanGTP inhibits cargo dissociation in animals but not yeast. **C)** RanGTP-bound ΔC loses its ability in cargo dissociation while yeast RanGTP-RanBP1 remains competent. **D)** ΔC dissociates cargo when its concentration is higher than that of RanGTP, and yRanBP1 dissociates cargo when its concentration is higher than that of CRM1.

Since ΔC-RanGTP did not bind to hCRM1 but still dissociated cargo (Fig. 1C and 2A), we therefore speculated that ΔC dissociates cargo through sequestering RanGTP from the export complex, which would result in a transient, low affinity binary complex of hCRM1 and NES cargo that automatically dissociates. Indeed, at high concentration of RanGTP, cargo dissociation activity of ΔC was fully inhibited (Fig. 2B), because excess of RanGTP not only saturated (and inhibited) mRanBP1, but also allowed CRM1 to form complex with GST-PKI. Further, purified 1:1 complex of ΔC-RanGTP, where RBD was already saturated with RanGTP, was unable to dissociate cargo (Fig. 2C). These results illustrate that ΔC, but not NES^mRanBP1^, is required for CRM1 cargo dissociation, through stripping RanGTP out of the nuclear export complexes.

### Cargo dissociation mechanisms in yeast and animals

We also performed the above cargo dissociation experiments using yeast protein in parallel. In contrast to mRanBP1, cargo dissociation ability of yRanBP1 was not inhibited by excess of Ran, and yRanBP1-yRan remained active to dissociate CRM1 cargo (Fig. 2B, 2C). Further, when Ran is in large molar excess than CRM1, yRanBP1 dissociated cargo as long as its concentration was higher than CRM1 concentration; in contrast, mRanBP1 dissociated cargo only at concentration that was higher than that of RanGTP (Fig. 2D). It is evident that yeast RanBP1 dissociates cargo through sequestering CRM1-RanGTP, distinctive from animal RanBP1’s cargo dissociation mechanism (Fig. 3A). Since CRM1 or more generally karyopherins, protect bound RanGTP from GAP mediated hydrolysis ^7,26^, the mechanistic difference of cargo dissociation predicts that hCRM1 would not protect RanGTP-RanBP1 from GAP hydrolysis like the yeast proteins. Indeed, the rate of hydrolysis in yeast was partially inhibited with addition of yCRM1, while there was no change of hydrolysis rate with addition of hCRM1 (Fig. 3B).

**Figure 3.**
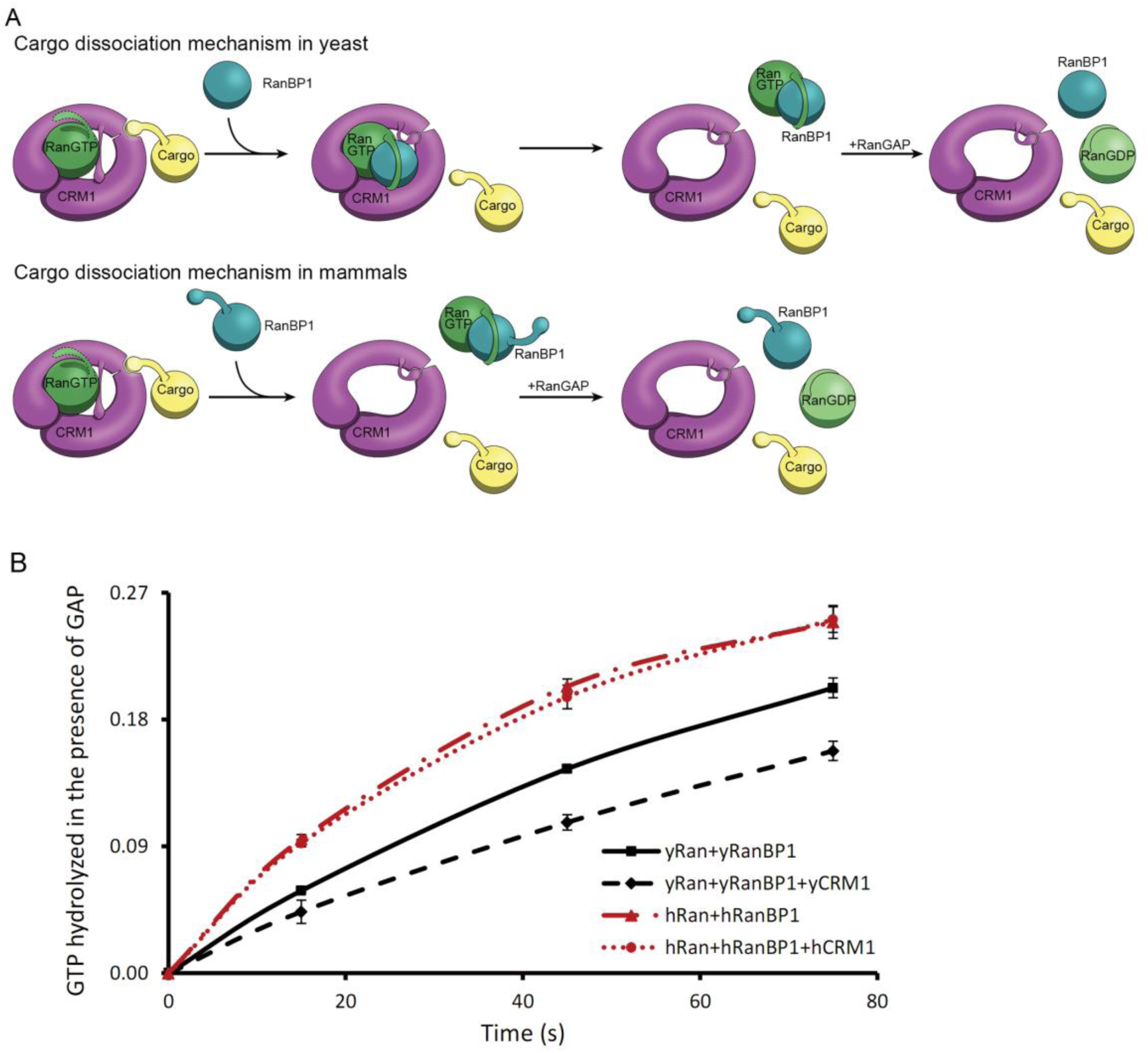
CRM1 cargo dissociation mechanism in fungi and animals. **A)** In fungi, cargoes are actively released from CRM1’s NES groove by formation of RanBP1- RanGTP-CRM1 trimeric complex. RanGTP-RanBP1 transiently released from CRM1 is catalyzed by RanGAP. In animals, RanBP1 strips RanGTP from CRM1, resulting in dissocaition of NES cargo from CRM1. RanGAP catalyzes further dissociation of Ran- RanBP1. **B)** GAP mediated RanGTP hydrolysis (arbitrary unit) in the presence or absence of CRM1. While addition of yCRM1 partially inhibit GAP mediated hydrolysis, addition of hCRM1 does not. Error bars represent standard deviation of quadruple repeats.

### RanBP1 forms complex with CRM1 and two RanGTP proteins in animals

Previously, we showed that mRanBP1 bound to hCRM1 through its NES and that ΔC did not dissociate cargo in excess of RanGTP. Since ΔC forms an extremely tight complex with RanGTP, we hypothesized that when RanGTP is excessive, mRanBP1 binds to one RanGTP though its RBD and binds to hCRM1 through its NES, while hCRM1 simultaneously binds to another RanGTP, forming a tetrameric complex (RanGTP-mRanBP1)-hCRM1-RanGTP (Fig. 4A). Unlike the yRanBP1 that contacts both yCRM1 and yRanGTP, animal RBD only bind RanGTP but not CRM1. In contrast to H9 loop stabilized closure of NES groove in yeast ^17^, animal CRM1’s H9 loop is far from NES groove, thus opening its NES groove for interaction with NES of RanBP1 (Fig. 4A). This model is consistent with all previous results, including that the binding between mRanBP1 and hCRM1 requires excessive RanGTP (Fig. 1A, B), that NES is the only interacting site between mRanBP1 and hCRM1 (Fig. 1C, D, F) and that ΔC-RanGTP does not dissociate CRM1 cargo (Fig. 2).

**Figure 4.**
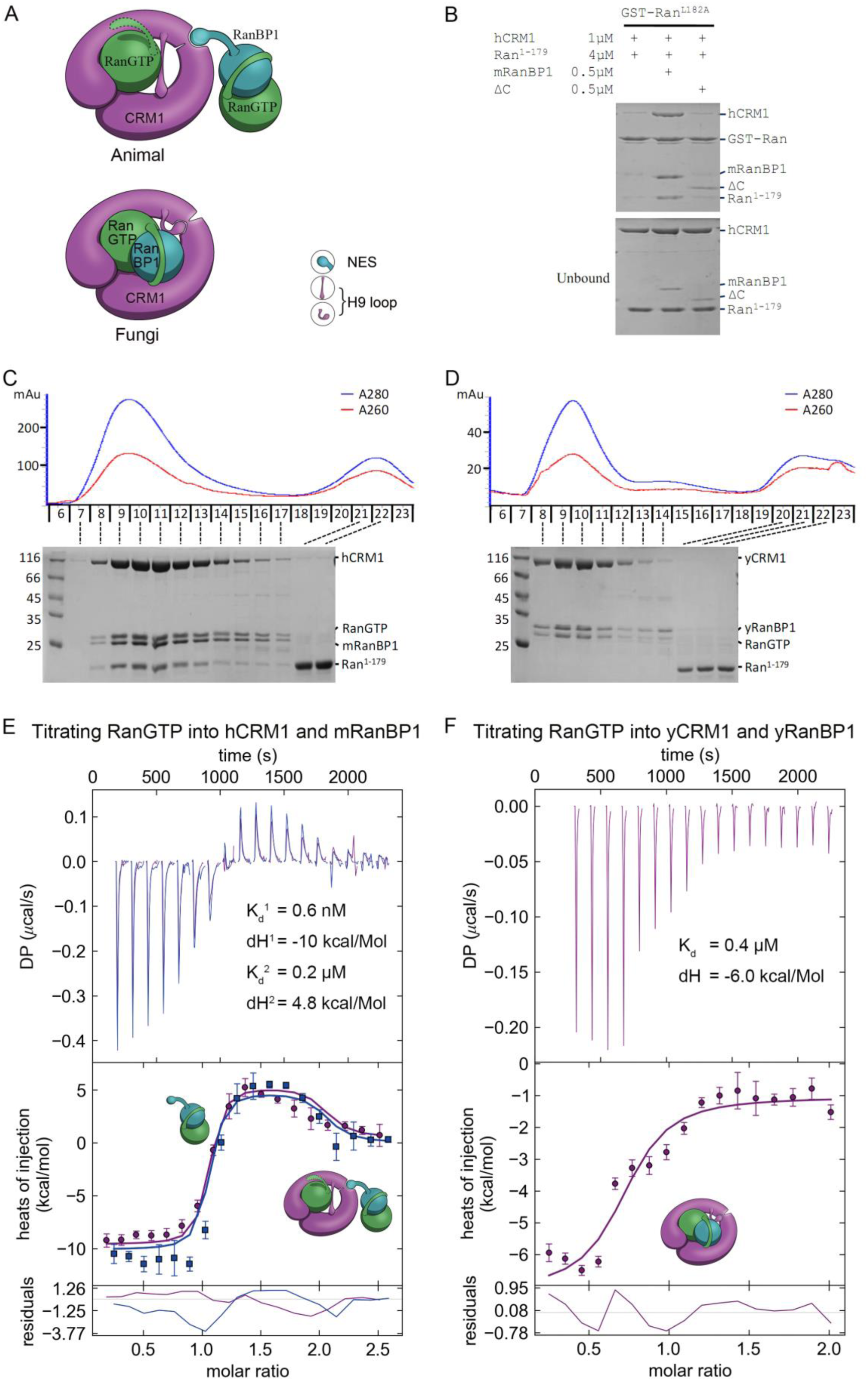
Animal RanBP1, RanGTP and CRM1 form a tetrameric complex containing two RanGTP proteins. **A)** Model of animal and fungi RanBP1 nuclear export complexes. Yeast complex contains one RanGTP and animal complex contains two. Unlike yeast, animal RBD does not contact CRM1 and animal CRM1 has different conformation of NES groove and H9 loop. **B)** GST-RanGTP pull down of Ran^1–179^ and CRM1 in the presence of mRanBP1, ΔC or buffer. Ran^1–179^ is bound when WT mRanBP1 is added, but not when ΔC is added. Size exclusion profiles and SDS-PAGE analysis of the peaks by animal **(C)** and yeast **(D)** RanBP1, RanGTP, RanGTP^1–179^ and CRM1 proteins mixed at 1:1:3:1 molar ratio. The first peak contains four bands for animal sample and three bands for yeast samples. ITC titration of RanGTP into animal **(E)** and yeast **(F)** CRM1-RanBP1 respectively. Figure E and F include two and one independent titration experiments respectively. Global or single fit of K_d_ and ΔH is displayed in the figure respectively. Error bars represents 95% confidence interval of measurements. While animal proteins display an exothermic phase and an endothermic phase, yeast proteins only produce an exothermic phase.

To further validate this model, we analysed the size exclusion peaks of yeast and animal RanBP1-RanGTP-CRM1 complexes mixed at 2:5:1 molar ratio. The calculated Ran to RanBP1 molar ratio in the peak of the animal complex was approximately twice (2.05 fold) than that of the yeast complex (Fig. S4), suggesting that animal complex contains two Ran proteins. To differentiate the two Ran proteins in the tetramer, we performed a pull down assay using GST-RanGTP and a truncated form of Ran, Ran^1–179^, which binds to CRM1 but does not bind to mRanBP1 due to lack of C-terminus. Clearly, Ran^1–179^ and hCRM1 bound to GST-RanGTP beads in the presence of mRanBP1 but not ΔC or buffer (Fig. 4B). The bound Ran^1–179^ can be explained by the formation of the proposed tetrameric complex, where GST-RanGTP binds to mRanBP1, which further binds to hCRM1-Ran^1–179^ through the NES of mRanBP1. Though ΔC binds to GST-RanGTP, the lack of NES rendered it unable to bind CRM1, therefore lack of Ran^1–179^ band in bound fraction. By size exclusion chromatography, hCRM1, RanGTP, mRanBP1 and Ran^1–179^ formed a reasonable 1:1:1:1 complex, whereas Ran^1–179^ did not co-elute with CRM1, RanGTP and RanBP1 in yeast (Fig. 4C, 4D). Further, RanGTP titration into hCRM1 and mRanBP1 by ITC produced an exothermic phase and an endothermic phase, representing the heat change from forming RanGTP-mRanBP1 and the tetrameric complex respectively (Fig. 4E). In contrast, the titration of RanGTP into yCRM1 and yRanBP1 displayed merely an exothermic phase representing the formation of the trimeric complex (Fig. 4F). Taken together, we conclude that RanBP1 forms a tetrameric complex with CRM1 and two RanGTP proteins in animals (when RanGTP excessive), where RanBP1-RanGTP binds to CRM1-RanGTP through the NES of RanBP1.

### Reasons for not forming trimeric RanBP1-RanGTP-CRM1 complex in animals

The key nuclear export and cargo dissociation mechanistic change from yeast to animals is the loss of binding between RanBP1-RanGTP and CRM1. We then asked why animal RanBP1- RanGTP did not bind to CRM1 as the yeast proteins. Through a pull down assay to test inter- species CRM1-RanGTP-RanBP1 complex formation, we found that in contrast to yRanBP1- hRanGTP-yCRM1 complex, neither yRanBP1-hRanGTP-hCRM1 nor ΔC-hRanGTP-yCRM1 complex was formed (Fig. 5A). Therefore, neither hCRM1 nor ΔC is compatible for trimeric complex formation, suggesting sequence divergence of essential residues (for trimeric complex formation) in yCRM1 and yRanBP1. We previously showed that yRanBP1 was unexpectedly localized in the nucleus when transfected into HeLa cells (Fig. 1F). This could be explained by Figure 5A since hCRM1 does not form trimeric or tetrameric complex with yRanBP1-RanGTP and thus is unable to export yRanBP1 into the cytoplasm in HeLa cells.

**Figure 5.**
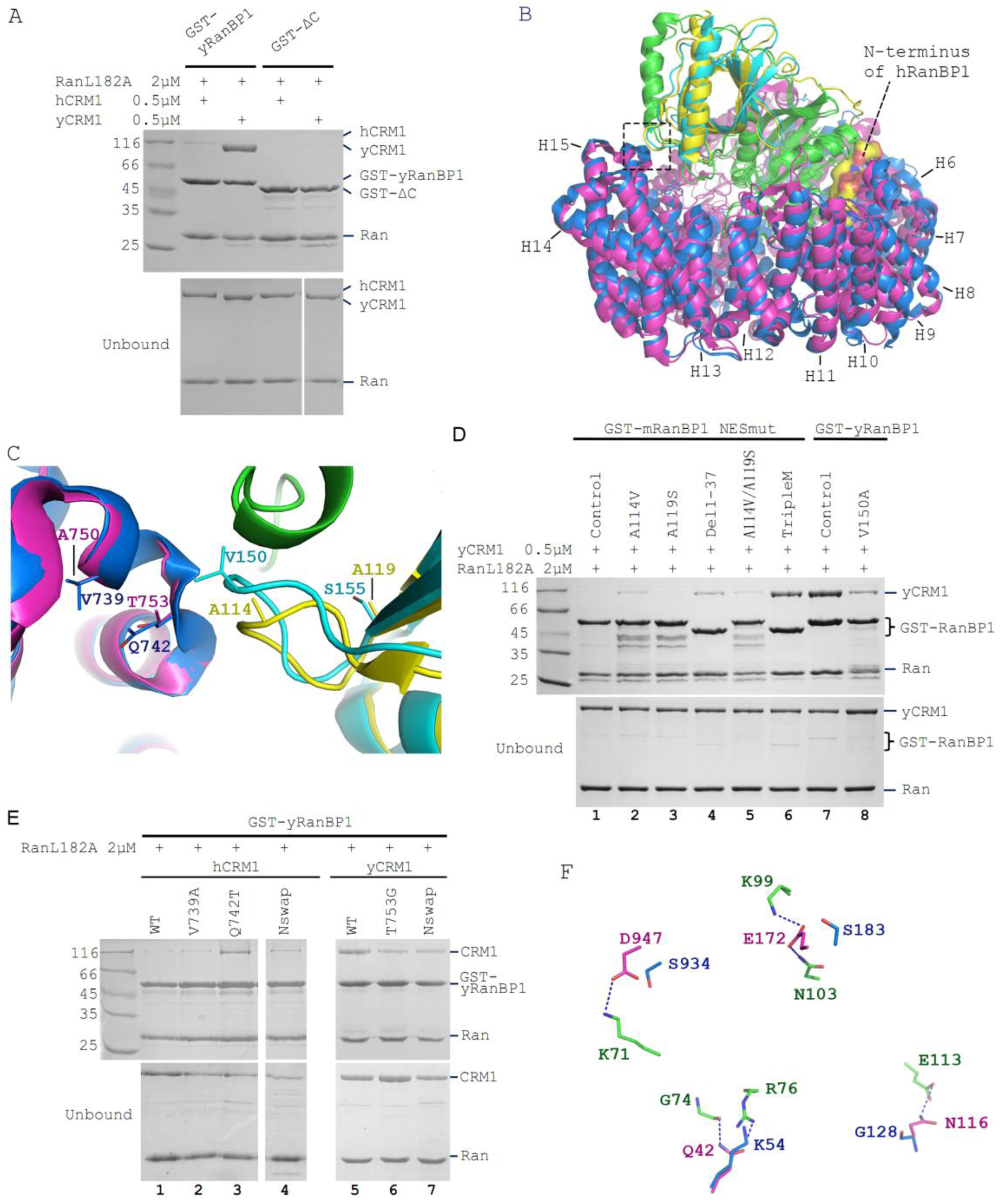
Sequence deviation of RanBP1 and CRM1 from their yeast orthologs prevents binding between RanBP1-RanGTP and CRM1 in animals. **A)** Cross-species pull down of CRM1 using immobilized GST-yRanBP1 or GST-ΔC in the presence of RanGTP. **B)** Superimposition of hRanBP1-hRan (1K5G, yellow and green) and hCRM1 (3GB8, blue) onto yRanBP1(cyan)-hRan(green)-yCRM1(magenta) crystal structure (4HAT). N-terminus of hRanBP1 (surface representation) is located between intimately packed RanGTP and CRM1 and thus possibly hinders their binding in animals. Heat repeats 6-15 (H6-H15) of CRM1 are labeled. **C)** Zoom in of boxed region in 5B shows RanBP1-CRM1 interface and adjacent residues (shown as sticks) that are different between species. **D)** GST-RanBP1 and its mutants pull down of yCRM1 and RanGTP. TripleM contains mutations A114V, A119S and Δ1-37, in addition to NES mutation. A114V and N-terminal deletion collectively rescue binding to yCRM1-RanGTP. **E)** GST-yRanBP1 pull down of human and yeast CRM1 mutants in the presence of RanGTP. Both RanBP1 and Ran interacting regions of CRM1 are important for trimeric complex formation. **F)** In the yRanBP1-hRan-yCRM1 crystal structure, yCRM1 and hRanGTP residues that form hydrogen bonds (dash lines) are displayed as magenta sticks. The structurally non-conserved hCRM1 residues are displayed as blue sticks. To improve clarity, main chain atoms (except glycine) and conserved residues are omitted.

When crystal structures of hRanBP1-hRan (pdb:1K5G) and hCRM1 (pdb:3GB8) were aligned onto yCRM1-hRan-yRanBP1 (pdb:4HAT), several differences between RanBP1 and CRM1 of the two species were observed (Fig. 5B,C). First, unlike yRanBP1 ^17^, hRanBP1 residues preceding its RBD domain are ordered and inserted between the closely packed Ran and CRM1 (Fig. 5B), possibly hindering trimeric complex formation by steric clash. Second, V150 that is in close proximity with yCRM1 in yRanBP1 corresponds to A114 in hRanBP1 (Fig. 5C). Interestingly, V150 is conserved to be valine or isoleucine in fungi (Fig. S5), yet not conserved in hRanBP2 and RanBP1 from human, mouse, zebrafish and frog (Fig. S2). The above discussed residues were mutated in NESmut and yRanBP1 to test whether they impact binding towards yCRM1 in the presence of RanGTP. Pull down assay results showed that either A114V or deletion of N terminus (Del 1-37) on NESmut could partially rescue binding to yCRM1. In agreement, V150A mutation in yRanBP1 partially inhibited binding of yCRM1 (Fig. 5D, lane 8 and 9). In contrast, mutation of adjacent residue A119 to yeast equivalent residue S in NESmut, did not significantly affect binding. Triple mutant (A114V, A119S and Del1-37) rescued yCRM1 binding to comparable level as yRanBP1 (Fig. 5D, lane 2- 7). Taken together, both the N-terminus and CRM1 interacting region of animal RanBP1 are incompatible for trimeric complex formation.

Similarly, several regions of CRM1 were mutated to identify critical sequence changes that abolished the formation of trimeric complex in animals. In the yRanBP1 binding interface, side chain of the strictly conserved yCRM1 T753 is in close contact with yRanBP1’s V150 hydrophobic side chain (Fig. 5C). The corresponding residue in human is Q742, being strictly conserved in animals. Pull down assay results showed that Q742T mutation partially rescued binding between hCRM1 and yRanBP1-RanGTP (Fig. 5E, lane 3). Similarly, T753G mutation in yCRM1 also reduced its binding to yRanBP1-RanGTP (Fig. 5E, lane 6). V739A mutation in hCRM1 (V739 euqivalent residues in yCRM1 is A), however, did not rescue binding (Fig. 5E, lane 2).

Besides the yRanBP1 interaction surface discussed above, yRanBP1 binds to yCRM1 indirectly through RanGTP. Human RanGTP forms 26 hydrogen bonds with yCRM1 in yRanBP1-hRan-yCRM1 structure (Fig. S6) and those residues that form hydrogen bonds are strictly conserved in yeast Ran (Fig. S7). However, six hydrogen bonds are predicted to be lost in human by sequence or structure based alignment of two CRM1 proteins (Fig. 5F, S8). Five hydrogen bonds are contributed by N-terminal residues (E172, Q42, N116) in yCRM1, and one by a C terminal residue (D947) (Fig. 5F, S6). Instead of making single mutations individually, the N-terminal 200 residues of hCRM1 was replaced with yeast equivalent residues 1-188 and the chimera protein (hCRM1^N_swap^) was able to weakly bind to yRanBP1- RanGTP (Fig. 5E, lane 4). Similarly, when the N-terminal 1-188 region of yCRM1 was replaced by human equivalent N-terminal region 1-200 (yCRM1^N_swap^), binding to yRanBP1-RanGTP was significantly reduced (Fig. 5E, lane 7). Pull down assay did not show a significant difference by further mutating D947S though (data not shown). Therefore, loss of binding between hCRM1 and yRanBP1-RanGTP is due to sequence divergence of hCRM1 from yCRM1, including the RanBP1 contact surface and Ran contact surfaces.

## Discussion

### Mechanistic difference between fungi and animal RanBP1 nuclear export

Nuclear export of RanBP1 is essential for all eukaryotic cells since excess of RanBP1 in the nucleus sequesters nuclear RanGTP, inhibits nuclear transport and is toxic to cells ^3,27^. In both yeast and human, RanBP1 is exported by nuclear export factor CRM1 ^19^. Though yeast and mammalian proteins are used in this work, the mechanisms revealed in this study should be applicable to fungi and animals. Due to the lack of NES, fungi RanBP1 should form trimeric complex through its RBD interacting with both RanGTP and CRM1. This explains the observation that intact RBD is required for its nuclear export ^19^. Obviously, NES is not needed for its nuclear export because fungi RBD is sufficient for its nuclear export. In animals, NES but not RBD is required for its export because animal RBD-RanGTP does not bind to CRM1. We show that this is due to degeneration of the interface residues essential for the formation of trimeric complex. Since nucleus is enriched with RanGTP, a tetrameric animal export complex of (RanBP1-RanGTP)-CRM1-RanGTP would form in animals, whereby RanBP1-RanGTP binds to the NES groove of CRM1 (through the NES of RanBP1) as an NES cargo. It should be noted that animal tetrameric complex probably exists only in the nucleus, whereas the fungi trimeric complex exists both in nucleus and cytoplasm. In the nucleus, animal RanBP1 cargo dissociation activity is inhibited with excessive RanGTP and it would not dissociate any CRM1 cargo or its own NES. When animal RanBP1 tetrameric complex enters the cytoplasm, where RanBP1 is present at high concentration and RanGTP at low concentration, cytoplasmic free RanBP1 binds to the RanGTP moiety bound to CRM1 in the tetrameric complex, accelerating dissociation of cargo (i.e., RanBP1-RanGTP complex).

### Mechanistic difference between fungi and animal RanBP1’s cargo dissociation

In the cytoplasm, fungi and animal RanBP1 also use different mechanisms to displace CRM1 cargoes. Both fungi and animals use RBD domains for cargo dissociation, and we showed that NES of animal RanBP1 is dispensable for cargo dissociation. Fungi RanBP1 binds tightly to RanGTP-CRM1 and closes NES groove to actively release bound NES. This ends up sequestering available CRM1, before RanGTP is hydrolysed with the help of RanGAP. In animals, excessive RanBP1 in the cytoplasm sequesters RanGTP from the NES-CRM1-RanGTP complex, resulting in a transient low affinity binary CRM1-NES complex, which dissembles automatically. However, we do not exclude the possibility that a transient complex of animal RanBP1-RanGTP-CRM1 forms like the fungi complex, which actively dissociates NES from CRM1, immediately followed by dissociation of RanBP1-RanGTP-CRM1. In fact, this is more likely since active displacement of cargo occurs with the yCRM1 mutant (P754D), which is defective in the stable yRanBP1-yRanGTP-yCRM1 ternary complex formation ^17^. Consistent with the proposed cargo dissociation mechanisms, we show that in the presence of RanGAP and RanBP1, CRM1 partially inhibits RanGTP hydrolysis in yeast but not in animals.

Besides dissociating cargo from CRM1, RanBP1 is essential for the disassembly of tightly bound Ran-karyopherin complexes. In fact, the Ran-sequestering mechanism seems to be more prevalent among RanBP1 mediated RanGTP dissociation from different karyopherins. Only human and yeast Importin β1 ^7,26,28,29^, and yeast CRM1 ^19,20^, form very tight complexes with RanBP1-RanGTP, which are partially GAP resistant. For human importin β2 ^7^, importin β3 ^28^, yeast importin 4 ^30^, human importin 7 ^31^, human importin 8 ^31^, human Exportin-1 ^8,32^, human Exportin-2 ^4,7^, human exportin-t ^33^, their complexes with RanGTP-RanBP1 are relatively weak and are effectively dissembled in the presence of RanGAP. In the case of human and yeast KPNB1, NLS is required for efficient release of Ran-RanBP1 for GAP hydrolysis ^7,29^. For yCRM1, there is no corresponding RanBP1-RanGTP release factor reported thus far.

### Biological significances of forming trimeric or tetrameric complex

Apparently, one round nuclear export of fungi RanBP1 costs energy of hydrolysing one GTP, while one round nuclear export of animal RanBP1 costs energy of two GTPs. Since RanBP1 is a constantly shuttling protein, animals may spend significantly more energy than fungi in the long term. We believe that at the same time of suffering this disadvantage in energy consumption, forming tetrameric complex may also confer at least two advantages for animals. First, as mentioned above, forming tetrameric complex does not inhibit GAP mediated hydrolysis, which means faster recycling rate of nuclear export machinery. This further implies possibly less congestion of nuclear pore and faster rate of nuclear transport. Second, the interface region and several other regions in fungi have to be conserved in order to maintain trimeric complex formation and export nuclear RanBP1. In animals, the existence of specialized NES in RanBP1 has eliminated that requirement, allowing multiple regions in both RanBP1 and CRM1 to diverge and participate in other yet-to-be discovered cellular functions. Compared with fungi, animals are readily mobile and more complex in cellular/tissue organization. Whether the appearance of RanBP1 NES in animals is involved in higher order cellular organization, tissue development or independent mobility warrants further studies.

Further, the different mechanisms observed in this study could aid in development of inhibitors against fungi by targeting the critical binding interfaces discussed above to inhibit the formation of trimeric RanBP1 nuclear export complex. Anti-fungal medicines developed by this strategy should be safe for human beings, since our RanBP1 nuclear export system does not require formation of trimeric complex.

## Conclusion

In summary, we have shown that animal RanBP1 binds to CRM1 only in the nucleus where there is excessive RanGTP, and forms a complex with CRM1 and two RanGTP proteins through it NES but not RBD. This complex is stable and is exported to the cytoplasm where it is dissembled by other RBD containing proteins and RanGAP. In contrast to CRM1-RanGTP sequestering cargo dissociation mechanism in fungi, animal RanBP1 dissociates cargo through stripping RanGTP from nuclear export complexes. The key difference between fungi and animals is that animal RanBP1-RanGTP does not bind to CRM1 as they do in fungi, due to degeneration of interface residues on both CRM1 and RanBP1, though either change is sufficient to abolish binding. The different mechanisms of RanBP1 nuclear export highlight how animals increase catalysis rate on the expense of more energy consumption. It is unexpected and interesting that while essential functions of orthologous proteins are conserved, profound differences exist in the underlining mechanisms.

## Acknowledgements

We thank Dr. Chong Chen, Dr. Yinglan Zhao and Dr. Bao Rui from Sichuan University for helpful discussions. Thanks Dr. Lei Qiu for proofreading this manuscript. This research is supported by Natural Science Foundation of China (NSFC) grants (#80502629, #31671477 and #81770580).

## Author Contributions

Q.Sun conceived the project. Y.Li, and J.Zhou performed biochemical and cellular works. Y.Zhang prepared several proteins used in this paper. Q.Sun, D.Jia and X.Shen supervised the project. Q.Sun prepared the manuscript. Q.Zhou and J.Han critically revised the article. The authors declare no conflict of interest.

